# Metaviromes reveal the dynamics of Pseudomonas host-specific phages cultured and uncultured by plaque assay

**DOI:** 10.1101/2021.04.27.441580

**Authors:** Katrine Wacenius Skov Alanin, Laura Milena Forero Junco, Jacob Bruun Jørgensen, Tue Kjærgaard Nielsen, Morten Arendt Rasmussen, Witold Kot, Lars Hestbjerg Hansen

## Abstract

Isolating single phages using plaque assays is a laborious and time-consuming process. Whether single isolated phages are the most lyse-effective, the most abundant in viromes, or the ones with highest ability to plaque on solid media is not well known. With the increasing accessibility of high-throughput sequencing, metaviromics is often used to describe viruses in envi-ronmental samples. By extracting and sequencing metaviromes from organic waste with and without exposure to a host-of-interest, we show a host-related phage community’s shift, as well as identify the most enriched phages. Moreover, we isolated plaque-forming single phages using the same virome-host matrix to observe how enrichments in liquid media corresponds to the metaviromic data. In this study, we observed a significant shift (p = 0.015) of the 47 identified putative *Pseudomonas* phages with a minimum 2-fold change above 0 in read abundance when adding a *Pseudomonas syringae* DC3000 host. Surprisingly, it appears that only two out of five plaque-forming phages from the same organic waste sample, targeting the *Pseudomonas* strain, was highly abundant in the metavirome, while the other three were almost absent despite host exposure. Lastly, our sequencing results highlights how long reads from Oxford Nanopore elevates the assembly quality of metaviromes, compared to short reads alone.

## 1. Introduction

As bacteriophages (phages) are the most abundant entity across all environments, they serve as a major reservoir of genetic diversity and have an essential roles in ecology, maintaining bacterial diversity as well as carbon and nutrient cycling [1–4]. Phages keep dominant bacteria in check by ‘killing the winner’ through lysis, which results in the re-lease of organic matter from the exterminated host used by other prokaryotes [5]. The ability of phages to attack and lyse very specific host-bacteria has sparked a significant interest to discover novel phages to fight pathogens in phage therapy (human infections) and phage biocontrol (agriculture) [6–11].

It is estimated that the number of phages is at least ten-fold larger than the number of bacteria in the sea and 1031 phage particles on earth [4,12–14]. The combined biomass of phages on earth is assumed to corresponds to 75 million blue whales [3,15,16]. Despite their importance and high number, phages remain understudied but research in phage genomics and viromics is on the rise. The number of available phage genomes in data-bases is rapidly increasing, especially the uncultivated viral genomes (UViGs), but there is a lack of high-quality phage genome representatives in the databases [14].

One of the reasons for the lack of high-quality phage genomes is that the methodology for isolating single phage strains with known hosts is laborious, time-consuming, and often the limiting step [17–19]. One of two methods (I) direct plating or (II) enrichment approach is most often used to isolates single phages. Enrichments are often used to increase the number of phages against the host-of-interest, and work by incubate the host together with a phage-containing sample in a growth media. Both ways are typically followed by double-agar overlay technique first described by Gratia to observe plaques forming units [20,21] An additional step used in some cases is polyethylene glycol (PEG) precipitation, which concentrates and purifies the phages in a sample before direct plating or after enrichment [22,23]. Plaque assays are usually the final step of isolating phage-host pairs. It is a simple technique, but parameters such as the nature of the phage itself, insufficient phage titer, temperature, percentage of the agar in the overlay, presence of certain ions or adsorption of phage particles to host cells can all affect whether or not a visible plaque will form [18,24]. Some phages are not capable of making plaques on plates due to limited diffusion in agar in general or low productivity, and isolation of highly effective single phages might be unsuccessful despite them being present and able to lyse the host [15]. All these restrictions are limiting types and numbers of phages isolated, despite attempts to make it more high-throughput [17].

Recent advances in sequencing has enabled genomic investigation of phages from environmental samples through targeted sequencing of metaviromes (referred to as viromes in this study) [25]. This allows for investigating both plaque-forming and non-plaque-forming phages, giving a broader understanding of the type of phages in different environments, and expand the phage-genomes databases.

Identification of non-plaque-forming phages targeting specific bacteria could improve the types of phages isolated against human-, animal- or agriculture-pathogens. Phages isolated by plaque assays may not be the most effective or the most abundant in the phage community, but simply the best ones to make detectable plaques. Thus, using a virome from an environmental sample, and extracting it with and without cultivation with a specific host, can highlight what members of the viral community amplifies in relation to the presence of the host in a complex viral community. One of the most important characteristics of a phage is describing which hosts it lyse or interact with. However, this characteristics is hard to predict based on the metavirome sequencing alone. Comparing the two types of viromes to phages isolated from plaque assays using the same host-virome matrix will help to predict phage-hosts further and indicate if the most abundant phages are plaque-forming, or whether other (perhaps non-plaquing) phages are enriched.

In this study, we use organic waste (OW) samples together with the tomato pathogen, *Pseudomonas syringae* pv. tomato DC3000 (DC3000) to analyze the community shift that occurs in the OW virome when exposed to the tomato pathogen. We compare the OW virome to five plaque-forming phages isolated by traditional plaque-assay from the same virome-host matrix and demonstrate that only two of the plaque forming phages are potentially among the most enriched in the virome. The setup of the experiment and key findings are depicted in Figure S1. This study raises important questions on how future phage isolation is to be done, how we can predict phage-host from metaviromes and demonstrate the importance of long read sequencing data when assembling viromes.

## 2. Materials and Methods

### 2.1 Bacterial host strain and organic waste sample preparation

*Pseudomonas syringae* pv. tomato DC3000 (DC3000) [26,27] was maintained on LB solid or liquid media [28] and incubated at room temperature (approx. 21°C and shaking at 200 rpm). The waste-management company HCS A/S (Glostrup, Denmark) provided the liquid generated from common household organic waste (OW) and was kept at −20°C until use, in order to stop all biological activity until thawing. To prepare the OW phage community for spiking/enrichment with DC3000 (explained in section 2.3), 300 ml of OW sample was centrifuged for 10 minutes at 10,000g, and the supernatant was filtered through PDVF syringe filters (0.45 μm, Merck Millipore, Darmstadt, Germany) to deplete from organic and bacterial matters from the sample. The filtered OW sample was kept at 4°C overnight.

### 2.2 Plaque assay and isolation of single phages

The five single phages (DrKristoffer, OtownIsak, SummerboyErik, GhostToast and Hovsa) from the OW sample were isolated using direct plating on 0.6% soft agar double overlay, the same way as the three phages described in Jørgensen et al., (2020) but using *P. syringae* pv. tomato DC3000 as host [11,29]. For DrKristoffer and OtownIsak the OW sample were PEG purified as described in section 2.3 before direct plating. SummerBoyErik was isolated from enriched OW samples: aliquots of the OW sample were enriched over-night with *P. syringae* DC3000 before direct plating as described previously [11,29,30]. GhostToast and Hovsa were both isolated with direct plating from the original OW sample. The phage DNA extraction was made as described by Jørgensen et al. (2020) and Carstens et al., (2019) [11,29]. Sequencing libraries for DrKristoffer, GhostToast, OtownIsak and SummerBoyErik were prepared using NEBNext® Ultra II FS library prep kit for Illumina (New England Biolabs, Ipswich, USA), while the sequencing library for Hovsa was prepared with Nextera® XT DNA Kit (Illumina, CA, US). The paired-end libraries were sequenced on the Illumina iSeq 100 platform (2×150 cycles). Adapter trimming and genome assembly was performed with CLC Genomics Workbench v. 12.0.03 with default settings.

### 2.3 Virome spike and extraction

The series of in-the-bottle infections, referred to as virome spike, were carried out in 200 ml LB media to which 4 ml of the filtered OW virome sample was added together with MgCl_2_ and CaCl_2_ to a final concentration of 10 mM, and 1 ml of the overnight DC3000 or sterile water in case of the non-spiked virome. Samples were incubated at room temperature with 200 rpm shaking for 8 hours. The spiked virome and the non-spiked virome samples had technical triplicates. A modified PEG precipitation of the protocol ‘Harvesting, Concentration, and Dialysis of Phage’ (www.phagesdb.org) was used for the initial concentration of the spiked virome, non-spiked virome, as well as 250 ml of filtered OW sample, from here referred to as the baseline virome. The viromes with the concentration of 10% polyethylene glycol 8000 (Millipore, 1546605) and 1M NaCl were left shaking at 200 rpm and 4°C overnight, followed by centrifugation for 10 min at 10,000 g and 4°C to pellet the PEG precipitate containing the phages and removing the supernatant. The PEG pellets were re-suspended in 12 ml SM buffer (100 mM NaCl, 8 mM MgSO4, 50 mM Tris-Cl) each and relocated in 50 ml falcon tubes. To ensure complete homogenization of the PEG pellet, the tubes were left shaking at 200 rpm and 4°C overnight. The samples were centrifuged at 4°C, 10,000g the next day to remove PEG and recover the supernatant now containing the phages. The supernatants were filtered through 0.45 μm PVDF filters (Merck Millipore, Darmstadt, Germany) for optimal removal of PEG and potential bacteria. All viromes were concentrated from 12 ml down to approximately 200 μl SM-buffer using the Amicon® Ultra-15 Centrifugal Filter Devices 100K (Merck Millipore, Darmstadt, Germany), following manufactures recommendations. Both the spiked and non-spiked viromes were in technical triplicates, and the baseline virome had no replicates. Seven viromes were prepared and sequenced in this study.

### 2.4 DNA extraction, purification, and library prep

DNA extraction was done by phenol:chloroform method [31]. Briefly, the phage samples were treated with 5U of DNase I (A&A Biotechnology, Gdynia, Poland) for 30 min at 37°C, followed by treatment with 0.5% (v/v) sodium dodecyl sulfate (SDS) and 6U of pro-teinase K (A&A Biotechnology, Gdynia, Poland) for 1h at 55°C. The samples had 200μl 3M NH3Ac added and were then treated with an equal amount of phenol:chloroform to re-move proteins and bacterial debris. The aqueous phase was transferred to a new tube and the DNA was precipitated standard EtOH precipitation [31]. The DNA pellet was dissolved in sterile MilliQ water. DNA concentration and purity were measured with Qubit 2.0 fluorometer (Life Technologies, Carlsbad, US) and NanoDrop spectrophotometer (Thermo Scientific, Waltham, US), respectively. Libraries from all seven viromes were built according to the manufacturer’s instructions using the Nextera® XT DNA Kit (Illumina, San Diego, US) and sequenced as paired-end reads on an Illumina NextSeq500 platform using the Mid Output Kit v2 (2×150 cycles). A barcoded Nanopore library was build for the baseline virome using Rapid Barcoding Sequencing kit (SQK-RBK004) and sequenced on the MinION platform using an R9.4 flow cell (Oxford Nanopore Technologies, Oxford, UK). The Ligation Sequencing kit (SQK-LSK109) was omitted to ensure inclusion of poten-tial circular phage genomes, which cannot have adaptors ligated. Sequencing and base-calling was done using MinKNOW and Guppy v3.2.6, respectively. The purity and the DNA concentration are shown in Table S2, together with the number of reads after quality control.

### 2.5 Quality control and assembly

The Mosaic workflow (https://github.com/lauramilena3/Mosaic) was used for quality control, assembly, identification of putative viral contigs, viral population clustering, and abundance table calculation.

In total, 12 assemblies were done: 1) baseline (Illumina only), 2) merged triplicates for DC3000 + (Illumina only), 3) DC3000 + A (Illumina only), 4) DC3000 + B (Illumina only), 5) DC3000 + C (Illumina only), 6) merged triplicates for DC3000 − (Illumina only), 7) DC3000 − A (Illumina only), 8) DC3000 − B (Illumina only), 9) DC3000 − C (Illumina only), 10) merged samples for baseline, DC3000 + triplicates and DC3000 − triplicates, assembled with Illumina and scaffolded with baseline Nanopore reads, 11) baseline long-read assembly polished with Illumina merged samples for baseline, DC3000 + triplicates and DC3000 − triplicates, 12) baseline Nanopore assembly polished with baseline Illumina reads. Assembly 1-9 used metaSPAdes, 10 with hybrid metaSPAdes and 11-12 with Canu (11 listed on Figure 1, 12 listed Table S3 and S4).

**Figure 1.**
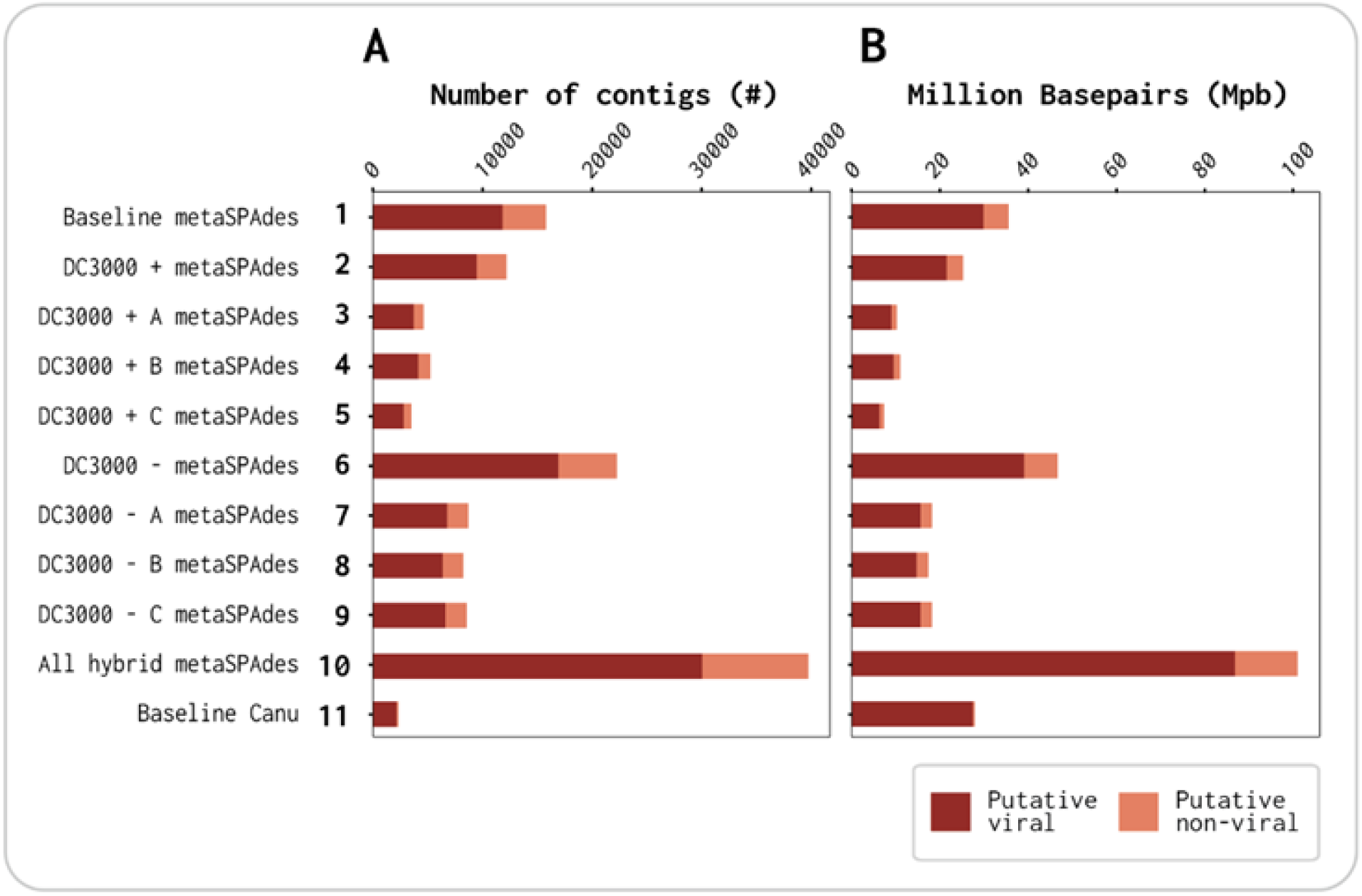
Identification of putative viral and putative non-viral contig count and Million base pairs (Mbp) in each of the 11 assemblies by VirSorter2 (A) Distribution of the number of putative viral vs. non-viral contigs. (B) Distribution of putative viral vs. non-viral Mbp.

For the quality control of Illumina reads, Trimmomatic 0.39 [32] was used to remove low-quality bases and adapters (LEADING:20 TRAILING:20 SLIDINGWINDOW:4:20 MINLEN:50). Trimmed reads were mapped with Kraken 2.1.1 against minikraken2_v2_8GB_201904 database to detect bacterial contaminants. Based on bacterial contaminants, reads mapping to ΦX174 (NC_001422.1), *P. syringae* pv. tomato str. DC3000 (NC_004578.1), plasmid pDC3000A (NC_004633.1), plasmid pDC3000B (NC_004632.1), *Lacticaseibacillus paracasei* (NC_014334.2) and plasmid plca36 (NC_011352.1) were removed with BBDuk from BBTools suite 38.86 [33]. Reads mapping to *Lactobacillus* are still present ranging from 10-11% in each of the seven viromes. Due to the lack of knowledge on the exact strain of Lactobacillus present in the OW, it was not possible to remove all Lactobacillus reads. Reads were normalized to ∼100x target coverage and were error corrected with bbnorm. For Nanopore reads, adapters were trimmed using Porechop 0.2.3 (https://github.com/rrwick/Porechop), and low-quality bases were removed using NanoFilt 2.7.1 [34]. Long-reads that mapped with minimap2 [35] to *P. syringae* or *L. paracasei* genomes were removed using SAMtools 1.10 [36].

For assemblies 1-9, down-sampled reads were assembled with metaSPAdes 3.14.0 [37] with no error correction. For assembly 9, down-sampled reads from each of the triplicates and trimmed Nanopore reads were used to perform a hybrid assembly with metaSPAdes. Assembly 10 of long-reads was done with Canu 2.0 (genomeSize=5m minReadLength=1000 corOutCoverage=10000 corMhapSensitivity=high corMinCover-age=0 redMemory=32 oeaMemory=32 batMemory=200) [38]. This assembly was error-corrected with long-reads using Racon 1.4.13 [39], and then polished with all the short-reads combined using Pilon 1.23 [40]. The resulting contigs from the 11 assemblies were concatenated and filtered to a minimum contig size of 1,000 bp and 2x coverage. Contigs denoted NODE_X are from metaSPAdes assemblies and contigs denoted tig0000X are from the Canu assembly.

### 2.6 Viral identification and clustering

VirSorter 2.0 [41] was run on all the assembled contigs to detect putative viral sequences. Identified viral sequences were merged with genomes of the five plaque-isolated phages and then clustered into viral populations (vOTUs, presumably clustered at species level) using stampede-clustergenomes script (https://bitbucket.org/MAVERICLab/stampede-clustergenomes/). Contigs were clustered at 95% nucleotide identity across 80% of the genome. The longest member of each cluster was selected as the representative. Completeness of representative contigs was assessed using CheckV 0.6.0 [42]. Contigs predicted as complete-quality are within the high-quality group. Viral population representatives were mapped to NCBI RefSeq bacterial genomes (downloaded on Feb 23, 2021) using BLAST (2.9.0+). Viral populations matching more than 40% of their length with a bacterial RefSeq hit were removed. Finally, vOTUs that which were predicted with CheckV as high-quality or medium-quality, and contigs less than 5,000 bp in those two groups were selected for abundance analysis too.

### 2.7 Abundance calculations

Quality controlled reads for the spiked and non-spiked replicates were subsampled to the lowest number of reads (2,228,236). Subsampled and total reads were mapped against the filtered viral clusters using BBMap 2.3.5 (https://sourceforge.net/projects/bbtools/). SAMtools [36] was used to generate a sorted bam. Contig depth coverage was calculated using tpmean from BamM 1.7.3 (https://github.com/Ecogenomics/BamM). In the same way, contig breadth coverage was calculated with genomecov from BEDtools 2.29.2.0 [43]. For a contig to be considered pre-sent, it was required that more than 80% of the sequence is covered. When this condition is not met, the abundance of the contig was set to zero. Viral population abundance tables for the total and subsampled reads were obtained, where the depth of coverage was normalized per million reads per sample. The relative abundance in each sample was calcuated by dividing the contig depth coverage by the sum of all contig depth coverages in that sample.

### 2.8 Host-prediction

SpacePHARER v4 [44] was used to predict phage-host relationships at genus level, based on the detection of vOTUs that matched to 363,460 CRISPR spacers from the Shmakov spacer dataset [45]. Alternatively, tblastx was used to align the vOTUs against a database of *Pseudomonas* phages. This database was obtained by downloading all 1,253 entries in the nucleotide archive of NCBI, which had “*Pseudomonas* phage” in their description. These entries were filtered and only 510 records that contained “complete genome” or “genome assembly” in their description were kept.

Contigs that matched a CRISPR spacer from a *Pseudomonas* host (SpacePHARER method), or contigs where more than 60% of the contig was covered by a sequence from the *Pseudomonas* phage database (tblastx method), were taken for manual validation. Those contigs were blasted using megablast, discontiguous megablast, or tblastx manu-ally to ensure validity [46]. To deem a contig to be a putative *Pseudomonas* phage, the top viral hit, with the lowest E-value and highest query coverage percentage, had to be a *Pseudomonas* phage.

### 2.9 Diversity Analysis and Statistics

Conda package scikit-bio 0.5.5 (http://scikit-bio.org) was used for diversity analysis.

Bray-Curtis dissimilarities between each sample were calculated, and the resulting distance matrix was used for ordination using Principal Coordinates Analysis (PCoA) and hierarchical clustering of the samples. Furthermore, analysis of similarities (ANOSIM) was used to test whether our distances within- and between category groups were different.

Differential abundance (DA) was done on the subsampled vOTU table using DAtest 2.7.17 [47], which through simulations, screens a variety of DA tests for ability to control both false negatives and false positives. MetagenomeSeq featuremodel [48] was selected, and used for analysis of differential abundance of vOTUs based on the sample being incubated with or without DC3000 (+/−). The log2 fold change (Log2FC) and False Discovery Rate (FDR) adjusted p-values were calculated by the MetagenomeSeq feature model, indicating the differential read abundance between the two types of viromes for all the included contigs.

To reveal if there was an enrichment of *Pseudomonas* phages in the spiked virome, a Fisher exact test was conducted on the DA results, comparing ratios of the curated putative *Pseudomonas* phages compared against the putative *Lactobacillococcus* phages detected with the CRISPR method. *Lactobacillus* phages were selected as reference due to its relative high coverage and since they are naturally present in a high abundance in the OW sample.

## 3. Results

### 3.1 Viral contigs constitute the majority of each assembly

A total of 12 assemblies were generated using either metaSPAdes (v. 3.14.0, assembly 1-9), hybrid metaSPAdes (v. 3.14.0, assembly 10), or Canu (v. 2.0, assembly 11-12) with the Illumina reads, Illumina and Nanopore reads, or Nanopore reads polished with Illumina reads, respectively. These 12 assemblies compose of the baseline (assembly 1), one assembly for each of the DC3000 + A-C viromes (assembly 2-4), as well as one assembly for the combined Illumina reads of DC3000 + (assembly 5). The last six assemblies are each of the DC3000 − A-C viromes (assembly 6-8), one of the combined reads Illumina of DC3000 – (assembly 9), one with all Illumina and Nanopore reads concatenated from the seven viromes (assembly 10). Finally, a baseline with the long reads for assembly and polished with all short reads (assembly 11) and a baseline with the long reads for assembly and polished with short reads from only the baseline (assembly 12). Assemblies 1-11 are listed in Figure 1 with their respective assembler and 1-12 are listed in both table S3 and S4.

The distribution of putative viral, non-viral contigs and bases were identified with VirSorter2 in all assemblies (Figure 1A,B and Table 1, and detailed data for individual assemblies in Table S3 and Table S4). This showed that >75% of the assembled contigs were identified as viral in each assembly (Figure 1A and Table S3), and of the total 173,915 contigs assembled, 58% or 101,068 were identified as putative viral (Table 1). The viral contigs consist of relatively long contigs as >84% of the bases are identified as viral in each assembly (Figure 1B, Table S4). The baseline Canu (assembly 11) has the fewest total contigs (2297), but is the assembly with the least amount of contamination of non-viral contigs (113) and bases (1.1% non-viral Mbp vs. 98.9% viral Mbp) (Table S3 and Table S4). These results highlight the importance of long reads, corrected with short reads, as the baseline metaSPAdes (assembly 1) using only Illumina reads has a total of 15,782 contigs, where 75.2% are putative viral and 24.8% of the Mbp are putative non-viral (Table S3 and Table S4). The amount of putative viral vs. non-viral Mbp correlates with the length of the contigs for the two baseline assemblies as N50 is 2,901 bp and 21,781 bp for the metaSPAdes (assembly 1) and Canu (assembly 11), respectively. Table S3 and S4 show that there is little to no change in the identified putative viral contigs and Mbp by VirSorter2 between assembly 11 and 12. Assembly 12 was added to compare only baseline Nanopore reads vs baseline Illumina reads, to ensure that the longer contigs and lesser non-viral Mbp were a result of the long reads and not of polishing with all short reads from the seven viromes. Only assembly 1-11 will be used for further analysis.

**Table 1.** Data of total contigs and total Mbp from the merged 11 assemblies. Putative viral and putative non-viral contigs and Mbp were identified by VirSorter2 and are visualized for each assembly in Figure 1. Data for individual assemblies’ contig and Mbp distribution is denoted in Table S3 and Table S4. Assembly 12 is excluded.

|  | No. of contigs | % |  | No. of bp | % |
| --- | --- | --- | --- | --- | --- |
| Total | 131,073 | 100 | Total | 317,732,011 | 100 |
| Putative viral | 101,068 | 77.1 | Putative viral | 275,137,398 | 86.6 |
| Putative non-viral | 30,005 | 22.9 | Putative non-viral | 42,594,613 | 13.4 |

Each of the 11 assemblies was clustered into viral populations as viral Operational Taxonomical Units (vOTUs) with 95% identity over 80% of the genome, presumably at the species level (Figure 2A, Table S5). Similarly, the total 101,068 VirSorter2 putative viral contigs from the 11 assemblies (Table 1) were clustered into 35,070 vOTUs (Figure 2B and Figure S1). The representative vOTUs of each viral population were put through CheckV (v. 0.6.0) to assess the contigs’ quality in terms of phage genomes’ completeness. Each sample’s vOTUs were classified as either putative High-quality, Medium-quality, Low-quality, or Not-determined (Figure 2).

**Figure 2.**
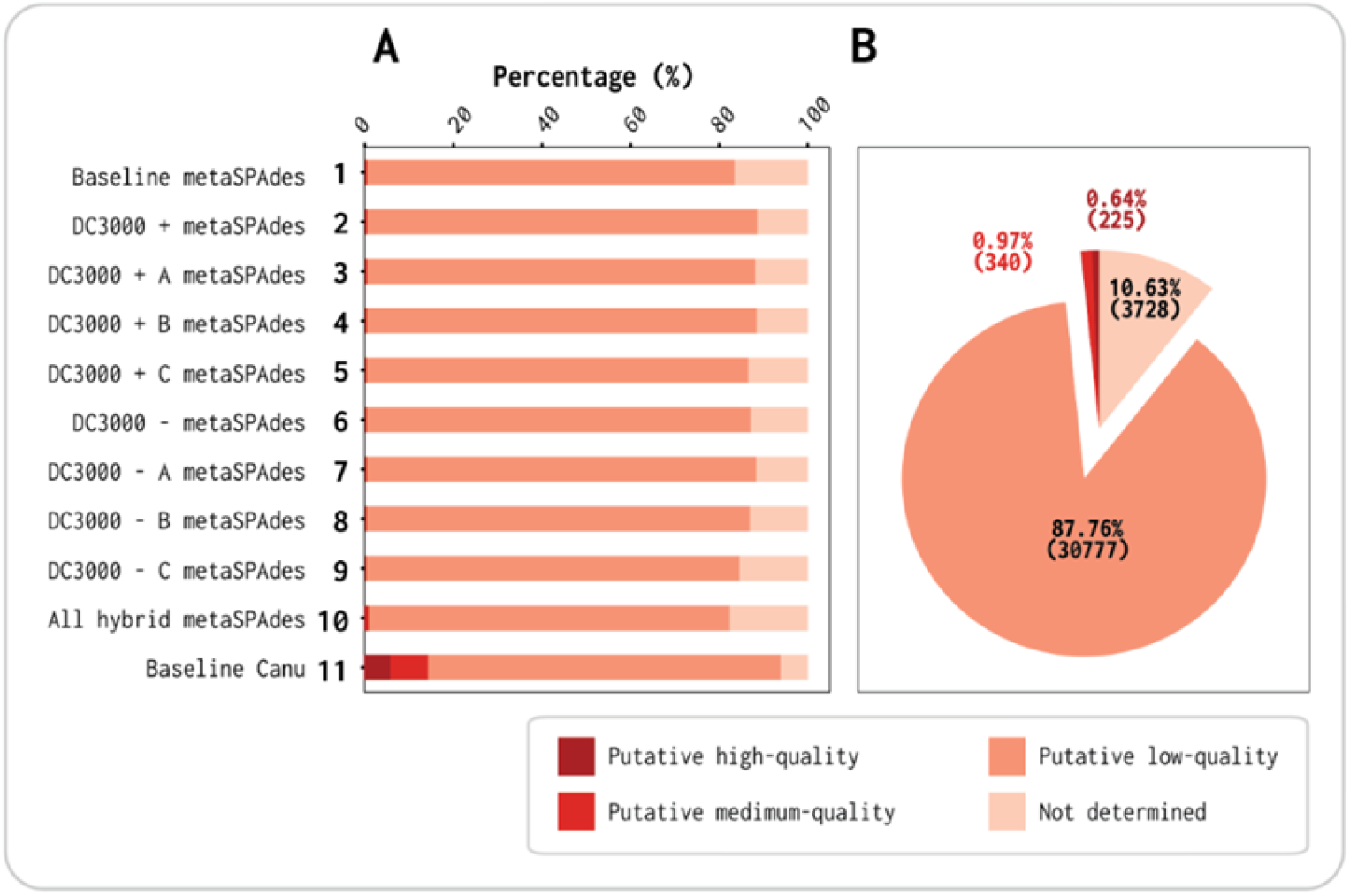
Quality identification by viral contigs with CheckV. **A** Percentage of quality distribution of viral contigs in each of the 11 assemblies. **B** Percentage of quality distribution of viral contigs for joined-assemblies.

The Canu assembler using the long reads and polished with the short reads gives a much greater amount of high-quality and medium-quality vOTUs than the assemblies using short reads only (Assembly 1-10 Figure 2A). The baseline Canu assembly (assembly 11) generated 2,004 vOTUs, while the baseline metaSPAdes (assembly 1) generated 14,788, which is close to 7 times more (Table S5). Of the 2,004 vOTUs from baseline Canu assembly, 116 were denoted as high-quality and 171 as medium-quality. Contrastingly, the baseline metaSPAdes (assembly 1) only had 29 high-quality and 51 medium-quality vOTUs despite the total contig number being 7 times higher (Table S5). Only by using all concatenated Illumina reads and scaffolding with Nanopore reads (assembly 10) more high-quality and medium-quality vOTUs were observed (132 and 196, respectively), but out of a total of 36,480 contigs (∼18 times higher contig no.) (Table S5). This means that the hybrid metaSPAdes (assembly 10) with Illumina and Nanopore reads from these seven viromes gave 0.36% high-quality vOTUs, while the Canu (assembly 11) resulted in 5.8%. This shows the difference in assembling with short reads and scaffolding with long reads vs. assembling with long reads and polish with short reads (assembly 10 vs. assembly 11).

When running CheckV on the total 35,070 vOTUs from this study, we see that the percentage of putative high-quality and medium-quality contigs is 0.64% and 0.97%, respectively (Figure 2B). These percentages would with high certainty been much higher if Nanopore sequencing had been possible for all seven viromes. Our results highlight (not surprisingly) how long reads raises the quality in terms of assembling phage genome completeness in the baseline virome (Assembly 1, 10 and 11). By using only or mostly short reads, many of the complete/high-quality viral contigs are possibly fragmented and predicted as low-quality.

Based on the Kraken 2.1.1 results, we observed reads taxonomically classified as bacterial, especially in the DC3000 + A, B, C viromes, potentially reflected in the difference of DNA concentration of said samples (Table S2). The bacterial genus with the most classified reads throughout all viromes was Lactobacillus. Furthermore, the three DC3000 + samples had reads mapping to the DC3000 strain too. Filtering steps to remove bacterial reads before assembly in all seven viromes are explained in section 2.5. Nanopore sequencing was feasible on the baseline OW virome, and a total of 12 Gb data was generated from the flow cell. The other six viromes did not extract enough DNA for Nanopore sequencing (Table S2).

### 3.2 Difference in viral composition correlates with presence/absence of vOTUs rather than high/low abundance

To remove false positive viral contigs post-VirSorter, all representative viral contigs were mapped against the NCBI RefSeq bacterial genomes database. All contigs covered > 40% to a bacterial RefSeq hit were removed. This ensured all contigs assigned to Pseudo-monas, Lactobacillus, or other bacterial species would be considered putative bacterial contamination and were not included for further analysis. All representative viral contigs < 5 kbp (unless they were denoted high-quality or medium-quality by CheckV) were removed for further analysis to avoid noise from small fragments. This resulted in a total of 3,515 vOTUs. Quality-controlled Illumina reads from the baseline, and each of the DC3000 + and DC3000 − viromes were down-sampled to the same number of reads. Reads from each of these seven samples were then mapped against the 3,515 filtered vOTUs to calculate the mean depth.

For each of the seven viromes, we observed 1,387 to 1,547 vOTUs (Table S8). To detect how similar their composition are to each other, we performed a hierarchical clustering based on the Bray-Curtis dissimilarities between each sample (Figure 3). We observed a difference between the viral population composition between the DC3000 + and − samples vs. the baseline. However, the DC3000 − had one sample (DC3000 − A) that clustered closer together with the DC3000 + viromes in regard to composition (Figure 3A,B,C). Changing beta-diversity to capture differences due to presence/absence (Jaccard) revealed similar results both in terms of hierarchical clustering and in relation to experimental design [49]. When comparing Jaccard and Bray-Curtis distance matrices with a mantel test we found a strong positive correlation (r=0.93, p-value=0.001). This indicates that the virome differences is driven by presence/absence of certain viruses and to a lesser extend their abundance (Figure S6). Thus, DC3000 − A could cluster together with the DC3000 + samples due to the stochasticity in which viral populations can evolve [50,51]. No bacterial host was added to this sample, only clean media and sterile water for the virome. Still, the two viromes DC3000 − B and C have a more similar viral composition to the baseline virome than any of the DC3000 + viromes (Figure 3A,B), and an overall difference between sampling groups is trending but not significant (ANOSIM test statistic = 0.58, p-value = 0.085, Figure 3D). Hence, the overall viral population composition is altered due to the addition of DC3000 to the sample, but is also affected by the incubation in growth media, as the difference is not significant between the virome sampling groups. The possibility for a non-bacterial free OW virome caused by the filtering step is discussed in section 4 as it can be a potential factor on how the overall composition of the viral population is altered.

**Figure 3.**
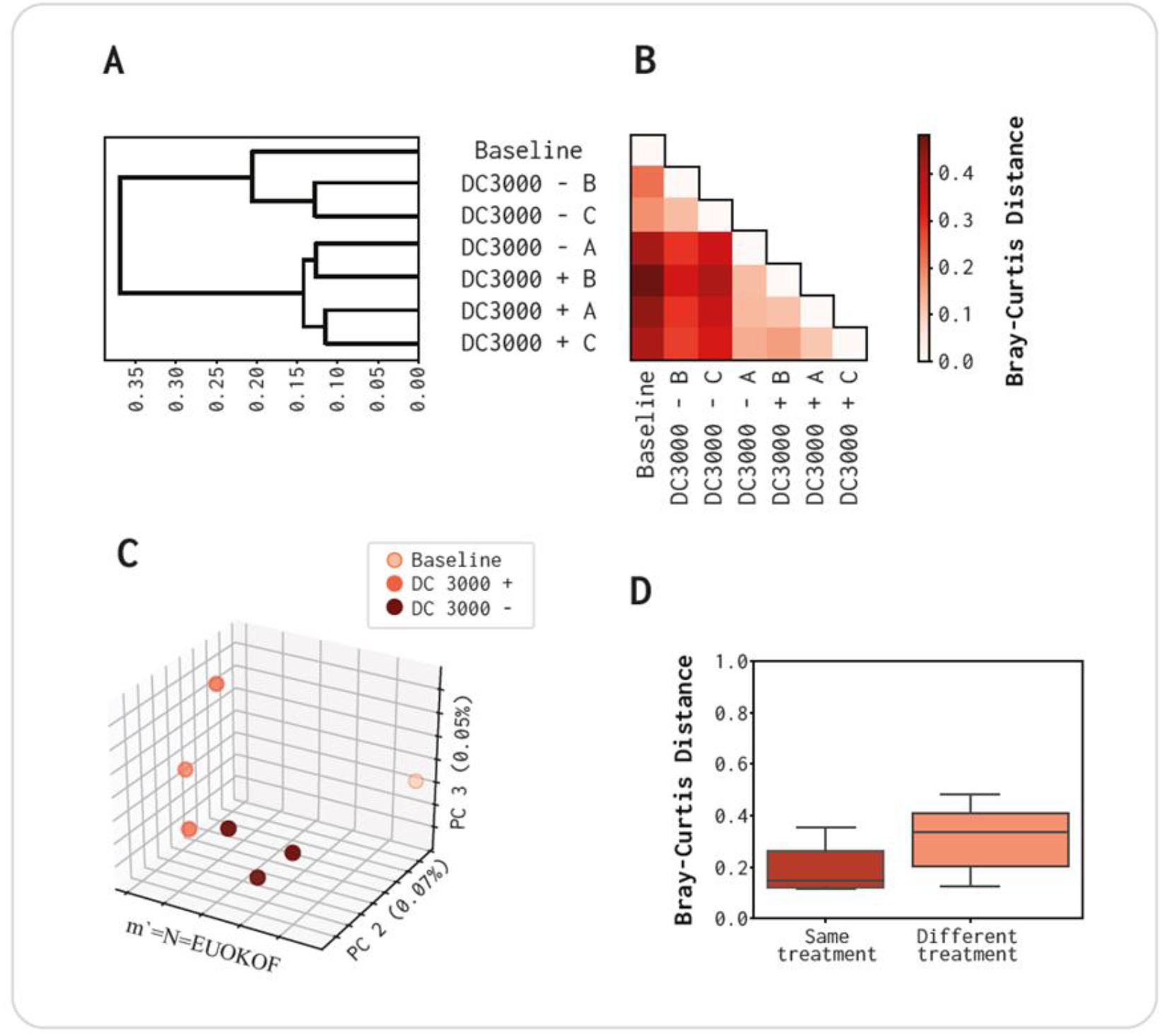
Viral OTUs composition of the population present in the three replicates of virome DC3000 +, DC3000 − and the baseline virome. A Hierarchical clustering of vOTUs based on Bray-Curtis distances. B Pairwise Bray-Curtis distances. C PCoA plot. D Comparison of treatment – incubation of OW in LB media at room temperature for 8 hours with or without an overnight culture of DC3000. ANOSIM test statistic = 0.58 (p-value = 0.085).

### 3.3 Most abundant vOTUs indicates a population change

The cumulative abundance indicated that the 10 most abundant contigs constituted ∼10% of the reads in the baseline virome, while they constituted ∼12.5% in the DC3000 + and DC3000 − viromes. Hence, the 10 most abundant vOTUs in the DC3000 + and DC3000 − account for around 2.5% more reads than the 10 most abundant contigs in the baseline assembly (Figure 4). A joined cumulative abundance graph of all contigs from the DC3000 +, DC3000 − and baseline metaSPAdes assemblies are denoted in Figure S7.

**Figure 4:**
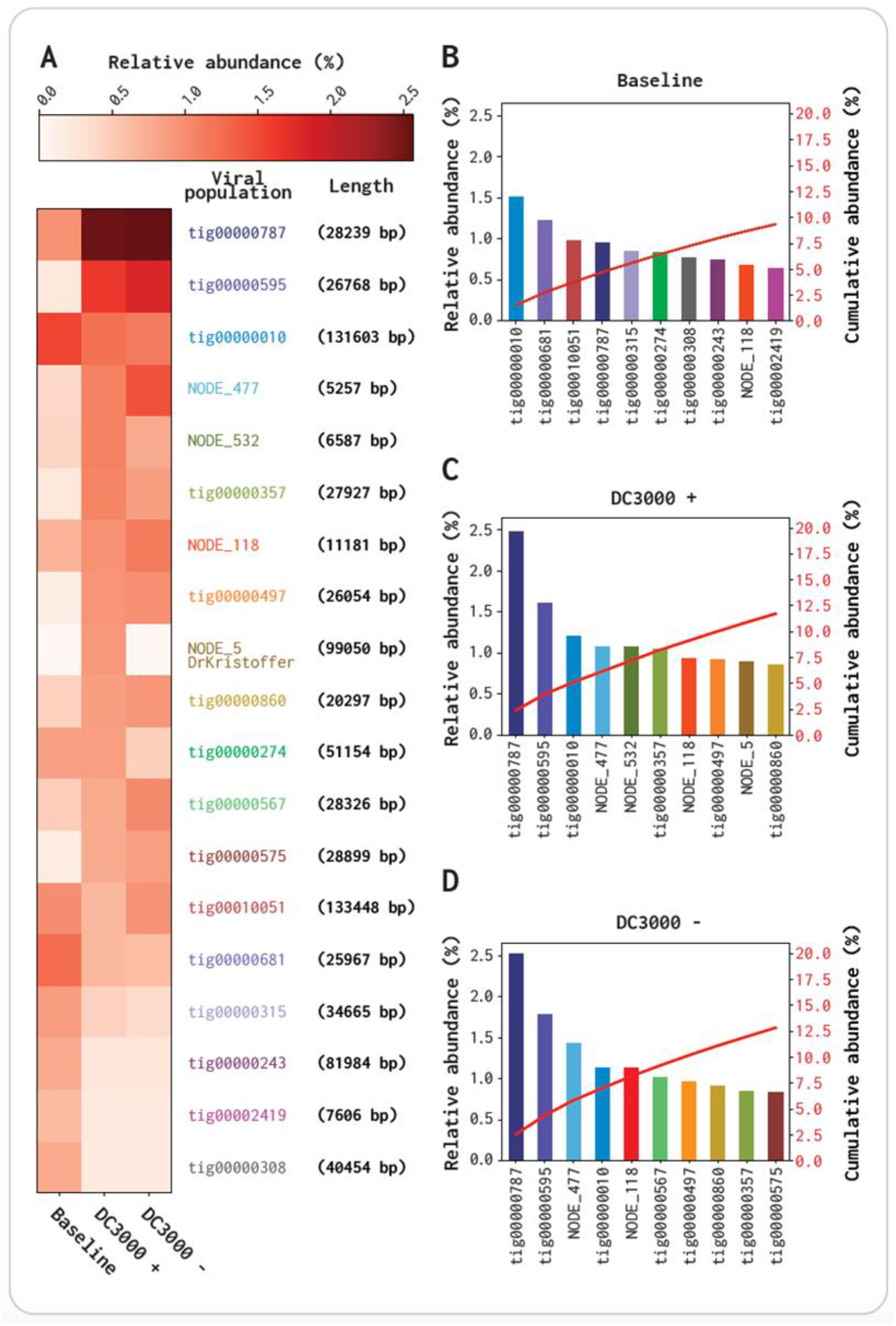
Abundance comparison of 10 most abundant contigs for the three viromes baseline, DC3000 + and DC3000 −. All contigs are putative viral identified by VirSorter2 and passed through the filtering and have < 40% query coverage to the bacterial database and is > 5 kbp. **A** Heat map of read abundance correspondent to the 10 contigs from each of the three viromes. B-D Relative abundance % (Left Y-axis), and Cumulative abundance % (Right Y-axis) of the 10 most abundant contigs in the baseline virome. **B** Baseline virome. **C** DC3000 + virome. **D** DC3000 − virome. It has to be noted that we did not account for whether these contigs could come from the same viral genome as fragments from the assembly.

The two contigs NODE_532 and NODE_5 are highly abundant in DC3000 + (1.04% and 0.87% relative abundance, respectively; Figure 4B,C,D). NODE_532 has lower read abundance in the baseline virome and DC3000 − (0.41% and 0.77%, respectively), and NODE_5 is completely absent in both samples (0.00% and 0.00%, respectively) (Figure 4A). A nucleotide blast alignment with the 99,050 bp long NODE_5 showed it has a query coverage and percent identity of 89% and 94.19%, respectively, and an e-value of 0 to the Pseudomonas phage VMC (Accession no. LN887844.1). NODE_5 is also observed to cluster together with one (DrKristoffer) of the five plaque isolated phages, which is further elaborated in section 3.4. NODE_532 is not absent in the baseline virome or DC3000 – (0.41% and 0.77% relative abundance, respectively) compared to NODE_5 (0.00% and 0.00% rela-tive abundance, respectively) (Figure 4A). A blast alignment showed it aligned to the Lactobacillus phage 3-521 with a query coverage and percent identity of 4% and 74.72%, respectively, and an e-value of 4e-41 (Accession no. NC_048753.1). As the query coverage is only 4% it is potentially only one gene that NODE_532 and the Lactobacillus phage 3-521 shares. In the group of 10 contigs most abundant contigs for each of the viromes, three (tig00000010, tig00000787, and NODE_118) are highly abundant in all three viromes, while five (tig00000595, NODE_477, tig00000357, tig00000497, and tig00000860) are highly abundant in DC3000 + and DC3000 − (Figure 4B,C,D). The baseline has seven contigs that are highly abundant, but are not within the group of 10 most abundant contigs in the DC3000 + and DC3000 − viromes. The two contigs, tig00000010 and tig00000787, are in the top four contigs in all three viromes, but they shift as tig00000010 relative abundance is reduced 0.37% in both DC3000 + and DC3000 −, whereas tig00000787 has a 1.4% higher depth coverage in both viromes compared to the baseline. The tig0000787 is also 0.85% and 0.75% higher in its accumulative abundance compared to the second most abundant contig, in DC3000 + and DC3000 −, respectively. These results indicate an abundance change happening potentially by the LB media incubation, as the shift is similar in both samples (Figure 4B,C,D).

### 3.4 DrKristoffer, Hovsa and OtownIsak cluster together with two contigs from the OW viromes

In parallel with the viromes, we isolated five single *P. syringae* phages from an aliquot of the OW sample also using the DC3000 strain as a host. These were included in the analysis to observe whether the isolated phages dominated the enriched population when the virome was subjected to encountering the DC3000. Their taxonomy at family and genus level is denoted in Table 2, together with their genome size, cluster representative and Blast comparison. All reads (not sup-sampled) were mapped to the five genomes and listed in Table S9.

**Table 2.**
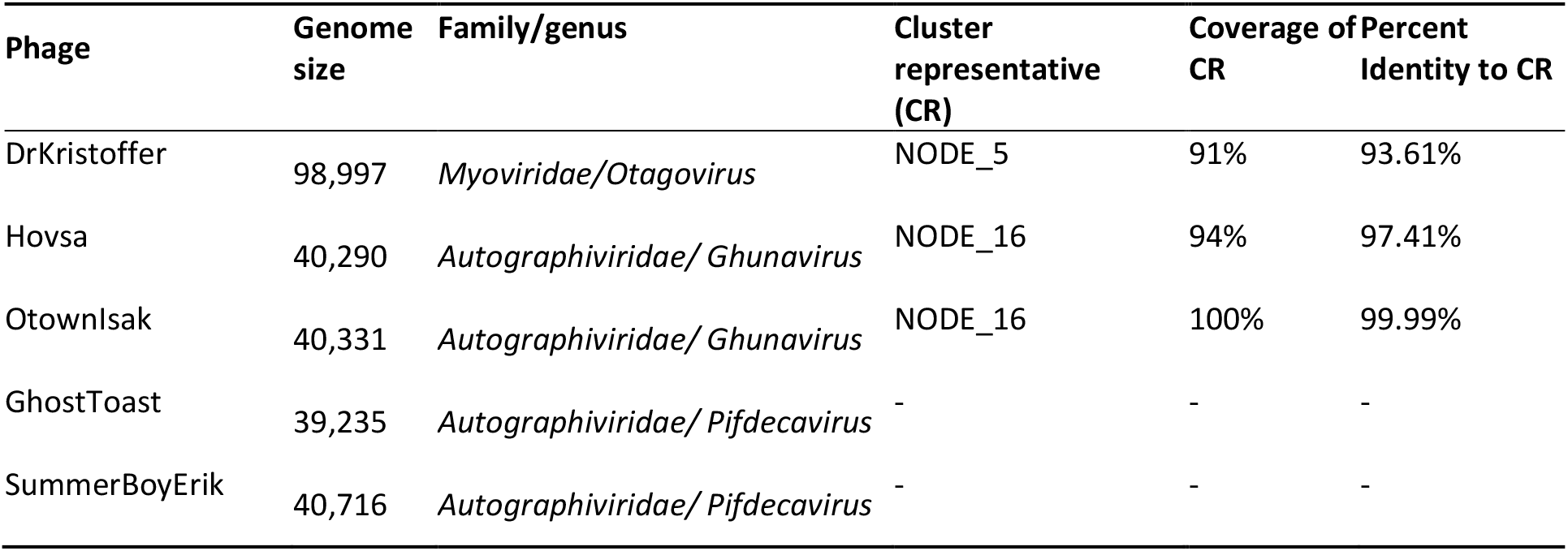
Overview of the five plaque-forming phages isolated against DC3000 from the same OW sample as used for DC3000 + and DC3000 − viromes. All blast hits had an E-value ≈ 0. GhostToast and SummerBoyErik did not have any Cluster representative.

Of the five phages that were able to make plaques on DC3000, only SummerBoyErik was incubated overnight with DC3000 in liquid media prior to plaque assay. GhostToast, Hovsa, and OtownIsak were all isolated with direct plating, whereas for the isolation of DrKristoffer, the OW sample was concentrated using PEG precipitation prior to plaque assay.

The genomes of the five phages were added to the analysis to observe which contigs they cluster together with (presumably at species level). NODE_5 (99,050 bp), which was mentioned in section 3.1 as one of the 10 most abundant contigs in DC3000 +. It is the representative of the vOTU cluster containing DrKristoffer and a blast alignment between NODE_5 and DrKristoffer showed a coverage and percent identity of 91% and 93.61%, respectively, and an e-value of 0 (Table 2). Hence, NODE_5 is potentially a phage from the same species as DrKristoffer. GhostToast and SummerBoyErik did not cluster to any putative viral contig in the viromes or to each other. The two phages Hovsa and OtownIsak, were members of the cluster represented by NODE_16 (40,386 bp). Due to the coverage and percent identity at 100% phage OtownIsak and NODE_16 could be considered the same phage (Table 2). However, Hovsa have a coverage and percent identity with NODE_16 at 94% and 97.41%, respectively (Table 2). These high values of both phages against NODE_16 means that we have to consider them, as the same species as NODE_16, but cannot state whether or not NODE_16 is OtownIsak or Hovsa. GhostToast and SummerBoyErik, are phages of the same family (*Autographiviridae*) as NODE_16/OtownIsak/Hovsa, but they are not related at species level to any contig in the viromes as DrKristoffer and NODE_16/OtownIsak/Hovsa. Mapping the total reads from each virome sample also against each individual phage genome, showed that DrKristof-fer/NODE_5 was only present in DC3000 + A, but NODE_16/OtownIsak/Hovsa was pre-sent in all seven viromes (Table S9). When comparing the isolated member of NODE_16 (OtownIsak and Hovsa) then OtownIsak had most reads mapping to its genome. This is, however, consistent with its ∼ 100% percent identity to NODE_16 and how BBmap classifies ambiguous reads according to the best scoring sites(Table 2 and Table S9) [52]. This could indicate that even though we isolated these phages from the same OW viromes, three of them (SummerBoyErik, GhostToast and DrKristoffer/NODE_5) are not highly abundant after incubation with DC3000 in all three replicates or in the baseline.

### 3.5 Pseudomonas phages are significant enriched when exposed to DC3000

From the 3,515 vOTUs (from section 3.2), 2,063 were present in at least one of the samples considering that at least 80% of the vOTU’s length needs to be covered for to be considered present. Hence, 2,063 vOTUs were included in the abundance comparison. Their p-values were plotted against the log2 fold change (Log2FC), where each dot represents contigs either getting more absent (Log2FC < 0) or present (Log2FC > 0) when the virome is incubated with the DC3000 host (Figure 5A).

**Figure 5.**
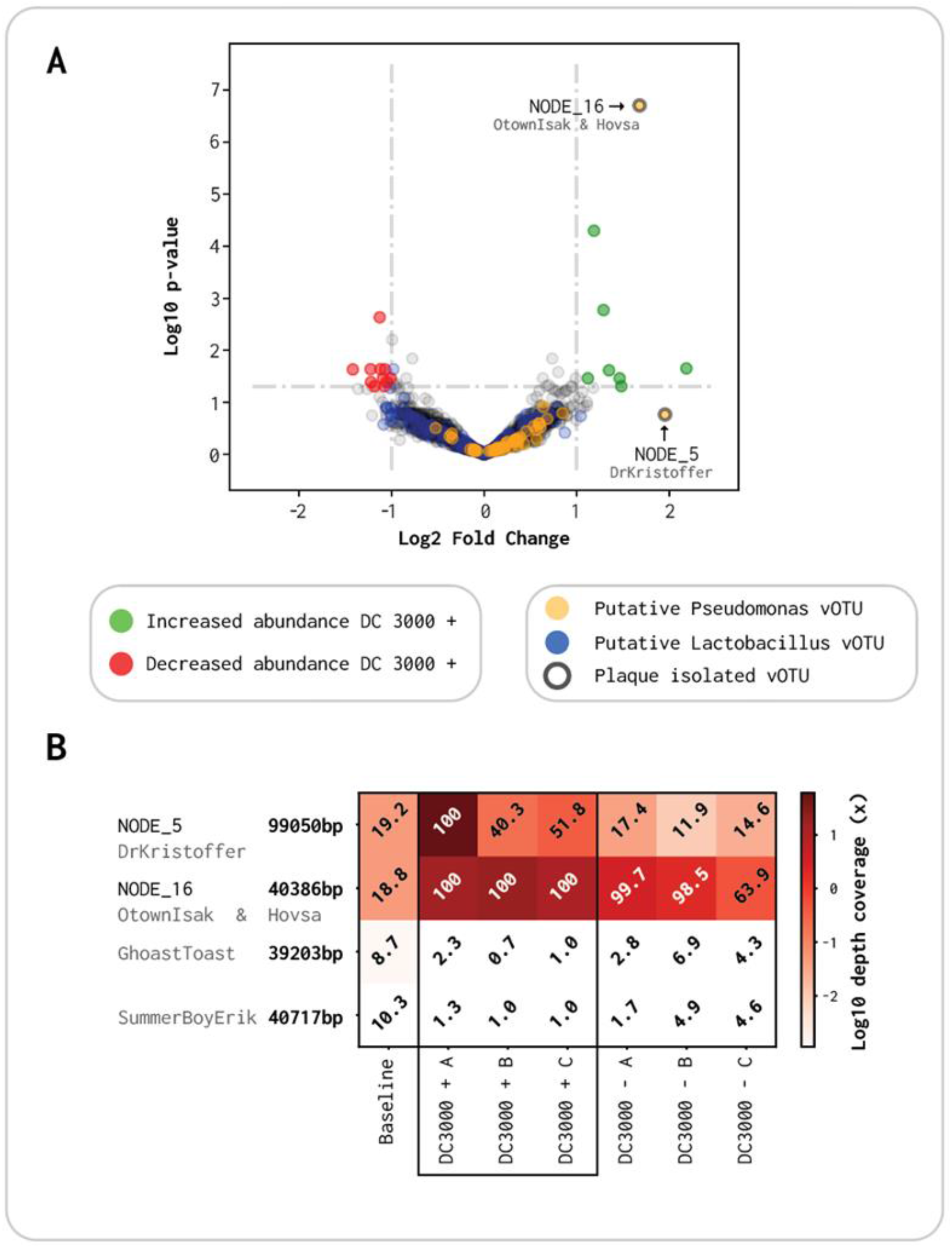
Volcano plot of abundance comparison between DC3000 + and DC3000 −. **A:** X-axis is log2 fold change (Log2FC) in read abundance between the two viromes, and Y-axis is log10 p-value for read abundance in the triplicates. Ringed dots: plaque forming phages Yellow dots: putative *Pseudomonas* phages. Green dots: viral contigs that are induced >1 Log2FC with a significant p-value. Red dots: viral contigs that are reduced <-1 Log2FC with a significant p-value. Blue dots: putative *Lactobacillus* phages **B:** Read abundance for the five plaque-forming phages throughout the seven viromes. White numbers indicate contigs with a breadth coverage >80%, which are presumed present, while black values indicate contigs with <80% breadth coverage regarded as not present.

A total of 8 contigs had a nominal significant p-value (α = 0.05) with a Log2FC above 1 (Green dots Figure 5A). These are the vOTUs enriched with the presence of the DC3000 strain. On the other hand, 11 contigs had a significant p-value with a Log2FC below −1, which are all phages getting reduced in abundance when the host is added (Red dots Figure 5A). The most abundant contig with a significant Log2FC of 1.68 is NODE_16, which we identified as the same species as OtownIsak and/or Hovsa in section 3.3 (Figure 5A and Table 2). It is the most significant contig (p < 2⍰10-7) and has a high read abundance in all three viromes of DC3000 + (Figure 5B). NODE_5 has a higher Log2FC of 1.95, but is not significant (p-value = 0.17). This is due to it being highly abundant in DC3000 + A, but not in B and C, and is therefore not significantly enriched compared to the DC3000 − viromes (Figure 5B). NODE_5 clustered together with DrKristoffer as they are the same species with a coverage of 91% (Figure 5 A and Table 2). The two last plaque isolated phages GhostToast and SummerBoyErik, had a very low abundance through all triplicates of DC3000 + and DC3000 −, compared to the three other single isolated phages (Figure 5B and Table S9). The seven additional contigs with a significant abundance and Log2FC > 1 in Figure 5 are all listed in Table S11, together with contigs with a significant abundance and Log2FC < −1. Contigs that had a significant abundance but a Log2FC between −1 and 1 are listed in Table S12.

Using CRISPR-spacer sequences analysis, we could assign hosts at the species level for a total of 503 out of the 2,063 vOTU. Of the 503, 32 (6.36%) were putatively assigned as Pseudomonas phages, and 209 (41.55%) as *Lactobacillus* phages. Alongside, the vOTUs were blasted against a database consisting of 510 *Pseudomonas* phages deposited in NCBI to further distinguish *Pseudomonas* phages. A total of 87 vOTUs were covered more than 60% with a contig from NCBI. The 32 CRISPR-spacer matching sequences and the 87 NCBI putative *Pseudomonas* (119 total) were manually curated as explained in section 2.9. Of the previous 119 vOTUs, 47 could be classified with high confidence as *Pseudomonas* phages, denoted as yellow dots (Figure 5A, Table S10). Moreover, 36 of the 47 (77%) curated *Pseudomonas* phages (Yellow dots Figure 5A) and 92 of the 209 (44%) CRISPR-spacer matching *Lactobacillus* phages (Blue dots Figure 5A) had a Log2FC > 0. We used Fisher’s Exact test to test whether the proportion of enriched *Pseudomonas* phages was significant revealing an odds ratio of 2.39 (Fisher’s Exact test CI: 1.16-5.30, p = 0.015), indicating a significant enrichment of *Pseudomonas* phages when incubated with a host organism, compared to *Lactobacillus* as a phage-host reference.

## 4. Discussion

When isolating single phages, the most used method is plaque assay [21,30]. Plaque-assay is also used to quantify recovered phages after virome extraction, when the virome sample is spiked with a known-phage in a high concentration to assess how efficient the virome extraction is [53]. To our knowledge, it is not well known whether plaque-isolated phages are the most responsive to the host-of-interest, or if they are simply the ones most capable of lysing the bacterial host in solid media.

Host-virus dynamics in liquid media has been analyzed before with di-lute-to-extinction isolation, but we hypothesized that when a virome is spiked in liquid media with a bacterial strain, novel to the phage community, there should be a shift in abundance towards phages targeting that strain [19,54]. Furthermore, we also hypothesized that the most enriched phages, in the spiked virome, are not necessarily those observed by plaque assay isolation. In this study, we mixed an overnight culture of *P. syringae* DC3000 with a filtered OW sample containing the virome in liquid LB media at room temperature for 8 hours. As a control, we added the same OW virome sample to liquid LB media without an overnight culture. By comparing these two types of viromes, we observed which phages are enriched by the *P. syringae* strain. Subsequently, we isolated five phages by traditional plaque assay from the same virome-host matrix. This allowed us to compare them to the most enriched vOTUs in the spiked virome. If they are highly abundant in the spiked virome, it will indicate that they are very effective in infection of the host in both types of media (solid and liquid). However, if they are low abundant it indicates out competition of these when mixed with other phages in liquid media and therefore have a much lower infection effectiveness when incubated with the host in solid media. Studying the shift of viromes when exposed to a specific bacterial host and analyzing how effective the single isolated phage(s) are in a different type of media will increase our general knowledge about phages and improve the methods we use to isolate phages that is non-plaque forming.

We assembled a large virome from OW with 2,063 putative vOTUs that is represented in at least one of the triplicates with a read coverage of >80% and could be used for reads abundance analysis. From this analysis we observed 8 putative phages that were significant upregulated with a Log2FC > 1 and 11 that were significant downregulated with a Log2FC < −1 (Figure 5A). Five dominant plaque-forming phages could be isolated from the OW sample using DC3000 as host, and the phages OtownIsak and/or Hovsa was indeed identified in the spiked virome as one of the most enriched vOTU to DC3000 (Table 2, Fig-ure 5A and Table S9). The phage DrKristoffer had a high Log2FC but was only highly abundant in one of the triplicates of DC3000 +, whereas the last two phages (Summerboy-Erik and GhostToast) had low abundance in all triplicates of both DC3000 – and + and did not have a breadth coverage above 80% (Table S9). This suggest that it is not necessarily the most abundant phages that are isolated by plaque assay. We show that comparative viromics can indicate significant host-specific dynamics, as we observed the proportion of enriched *Pseudomonas* phages was significant despite a large background of mainly *Lactobacillus* phages (Figure 5A).

A total of 3,515 putative vOTUs was identified across all seven viromes (Table S8 shows the amount of vOTUs in the individual samples). When comparing the vOTUs composition in the individual viromes, we found a strong positive correlation between the Bray-Curtis and Jaccard distance matrices (r=0.93, p-value=0.001) suggesting that the clustering is due to the presence/absence of vOTUs between the samples rather than abundance correlated. We observed that the composition of DC3000 − A was very similar to the DC3000 + triplicates (Figure 3). As mentioned previously, we filtered the samples with 0.45 μm PVDF filters to deplete from bacteria, but this filter size is not small enough to remove all bacteria, yet large enough to include larger phages otherwise lost with a pore size of 0.22 μm [55–59]. If the bacterial contamination is similar to the *Pseudomonas* DC3000 in any way, the sample can potentially evolve towards a similar composition as the DC3000 + viromes [55]. As the DC3000 − B and C more closely resemble the baseline virome, we assume that it is due to a stochastic variable, such as random bacteria passing through the filter, affecting the DC3000 – A (Figure 3A). The choice of a larger pore-size could potentially also be related to the 10-11% reads mapping to *Lactobacillus* in all seven viromes, as we see that DC3000 + and DC3000 − have a more similar shift in vOTU composition compared to the baseline (Figure 3C). If the OW sample had been completely removed of all bacteria by filtration it would be more likely, that the DC3000 − triplicates would be closer to the baseline in vOTU composition.

A total of 209 *Lactobacillus* phages we identified with CRISPR host-prediction in the 2,063 vOTUs, where the second most present genera are *Pseudomonas* phages with 47 (Ta-ble S10). The OW sample has a low pH value (≤ 3 pH) and as Lactobacilli prefers low pH value and often are associated with plant fermentation, it is likely that it is one of the dominant genera in the bacterial community [60,61]. To test if the proportion of enriched *Pseudomonas* phages was significant in the DC3000 + viromes, we used the Fisher’s Exact test. This showed that even though those 36 *Pseudomonas* phages did not significantly change in abundance, the proportion of them that increased was significantly higher than the ones that decreased. There were in average 2.39 more putative *Pseudomonas* phages with increased abundance in the DC3000 + viromes (95% CI, [1.16, 5.30]). Hence, we have shown a significant host-specific shift in the OW viromes when incubated in liquid media together with the DC3000 strain.

A total of 14 vOTUs had a significant difference in read abundance but a Log2FC be-tween 1 and −1, of which none were aligning to *Pseudomonas* phages, but six were similar to *Lactobacillus* phages (Table 10S). Additionally, 11 contigs were significantly different in their read abundance with a Log2FC < −1, of which four were similar to *Lactobacillus* phages and no *Pseudomonas* phages (Table 11S). One of those four contigs was the complete genome of the Lactobacillus phage 3-521 (accession no. NC_048753.1) isolated from Irish wastewater [62].

Eight contigs had a highly significant p-value (α = 0.05) with a Log2FC above 1 (Table S11). Three of those were putative *Lactobacillus* phages, while one (NODE_3) was the com-plete genome of the Enterobacteria phage CAjan (accession no. KP064094.1) isolated from rat feces [63]. The most significantly abundant of those eight was NODE_16, which is potentially the complete genome of the phage OtownIsak or/and Hovsa (Figure 5A). Both phages are isolated with plaque assay as described earlier together with GhostToast, SummerboyErik, and DrKristoffer. The phage DrKristoffer is the same species as NODE_5, which has a higher Log2FC than NODE_16 but with an insignificant p-value due to it having a high read abundance in only one of the DC3000 + viromes. Both GhostToast and SummerboyErik had a low abundance in all triplicates of DC3000 + and DC3000 −. If all five phages were the most effective of infecting and lysing the DC3000 strain in liquid media, their abundance would have been significantly high in all DC3000 + triplicates. As only OtownIsak or Hovsa can be identified in a significant read abundance, it is plausible to assume that the three other phages are not effective in lysing in liquid media but are able to do lysis in solid media as they can form plaques. Surprisingly, SummerboyErik was isolated by enrichment in liquid LB before direct plating with soft agar double overlay. From this, we would have expected SummerboyErik to be highly abundant after 8 hours, but perhaps the phage has a longer lag-phase and would have been significantly abundant in the DC3000 + viromes if they were left for longer. Another reason could be a limited burst size that would, which would mask the abundant of SummerBoyErik to the highly abundant phages.

Extracting the virome directly from the OW sample as the baseline virome, we were able to acquire long reads that included, in theory, all the initial phages present in the OW sample. The DNA concentration from the DC3000 + and DC3000 − triplicates were not sufficient for Nanopore sequencing (Table S2). Multiple displacement amplification (MDA) was not an option for this study as it has been shown to introduce bias such as over-amplification of small circular genomes and under-amplification of DNA with high GC content in viromes from human saliva [64]. The comparison of virome quality from the metaSPAdes assemblies and the Canu assembly further supports the importance of long-read data when performing metaviromics projects (Figure 1 and 4). Even though the baseline metaSPAdes (assembly 1) has more vOTUs (14,788) than the baseline Canu (assembly 11 with 2,004), the N50 (metaSPAdes 2,343 bp, Canu 21,775 bp) and in the amount of high-quality vOTUs (metaSPAdes 29, Canu 116) is still greater for the Canu assembly. Furthermore, 95.7% of the length of the Canu assembled contigs are covered by Illumina reads, showing that the information obtained from both platforms was similar. Thus, this difference is possibly due to the fragmentation because of the presence of microdiversity on the viral popilations that cannot be resolved with only short reads [65]. When adding the long reads, we see an increase from 0.18% to 5.79% of high-quality phage genomes compared to using only short reads despite the higher difference in read amount (Table S2, Figure 2 and Table S5). For studies not doing comparative analysis as our study, amplification is possible. Oxford Nanopore Technologies do offer library kits with low input DNA requirement (Rapid PCR Barcoding Kit SQK-RPB004), but it still not close to what is possible with the low input DNA Illumina requires (Nextera® XT DNA kit) [25,66]. Viromes sequenced with a combination of long and short reads will elevate the quality of UViGs in the public databases compared to only using short reeds (Figure 2) [67]. Better databases with high-quality phage genomes would also raise the quality of the available bioinformatics tools for phage identification in metavirome data. We attempted to assemble the seven viromes before removing the bacterial contamination to investigate how VirSorter2 and CheckV would assess the assembled contigs and if the former would be able to remove the bacterial contigs [41,42]. Both tools identify contigs that, when blasted had 100% alignment to bacterial genomes and none to phage genomes. Hence, the harsh removal of all putative bacterial reads, though we lost potential data from prophages that we had observed before the filtering.

## 5. Conclusion

Our results show that the DC3000 + and DC3000 − viromes can reveal significant host-specific dynamics of the phages when incubating for 8 hours in liquid media. We al-so showed that three out of five phages isolated individually from the same sample virome with plaque assay are ineffective of lysis in liquid media or outcompeted by phages from the same virome. This has to be considered when deciding if the single isolated phages are good potential candidates for treatment in phage therapy or biocontrol [17,29,68–70]. Furthermore, our analysis highlighted how much long-read sequencing data improves the assembly of viral metagenomes as we saw an increase of 5.64% in the proportion of complete phage genomes from long-read assembly compared to short-read-only assemblies (p-value < 0.0001, Figure 2).

## Author Contributions

Conceptualization, L.H.H., W.K and K.W.S.A.; methodology, K.W.S.A., L.M.F.J., and M.A.R.; software, L.M.F.J. and M.A.R.; formal analysis, K.W.S.A. and L.M.F.J.; inves-tigation, K.W.S.A. and L.M.F.J.; data curation, K.W.S.A., L.M.F.J., T.K.N. and J.B.J.; writ-ing—original draft preparation, K.W.S.A. and L.M.F.J.; writing—review and editing, L.H.H., W.K., J.B.J., T.K.N. and M.A.R.; visualization, L.M.F.J.; supervision, L.H.H. and W.K.,; project ad-ministration, K.W.S.A.; funding acquisition, L.H.H and W.K. All authors have read and agreed to the published version of the manuscript.

## Funding

This research was funded by Human Frontier Science Program with grant number HFSP – RGP0024/2018 for K.W.S.A, and Novo Nordisk Foundation with grant number NNF19SA0059348 (The MATRIX) for L.M.F.J and L.H.H. This project has received funding from the European Union’s Horizon 2020 research and innovation program under the Marie Skłodowska-Curie grant agreement No 801199.

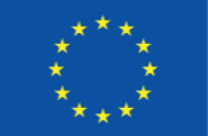

## Data Availability Statement

All reads used in this study are deposited at SRA and published under the BioProject ID PRJNA724013

## Acknowledgments

We would like to thanks Dr. Frederik Cold, who provided the names of the five isolated phages.

