## Supplementary material for "Metaviromes reveal the dynamics of Pseudomonas host-specific phages cultured and uncultured by plaque assay": Figure S1

**Figure S1:** Overview of the experimental set-up, sequencing platforms, and analysis. **Plaque assay:** Five single phages were isolated with either direct plating (Hovsa and GhostToast), PEG purification followed by direct plating (DrKristoffer and OtownIsak), or enrichment overnight similar to the incubation for DC3000 + and DC3000 – (SummerBoyErik). **Virome:** Visualization of the three types of viromes, DC3000 +, DC3000 – and Baseline. Incubation time for DC3000 + and DC3000 – was 8 hours at room temperature. **Keyfindings:** A total of 101,068 putative viral contigs was identified from all merged reads, and clustered into 35,070 viral populations (vOTUs), where 3,515 were >10 kb. We identified three of the five phages in the viromes, but observed that OtownIsak or Hovsa was the most enriched phage when incubated with DC3000 as they belonged to the same vOTU. Pseudomonas phages were confirmed to be significantly enriched when incubated with DC3000. Created with BioRender.com and restructured in Adobe Illustrator.

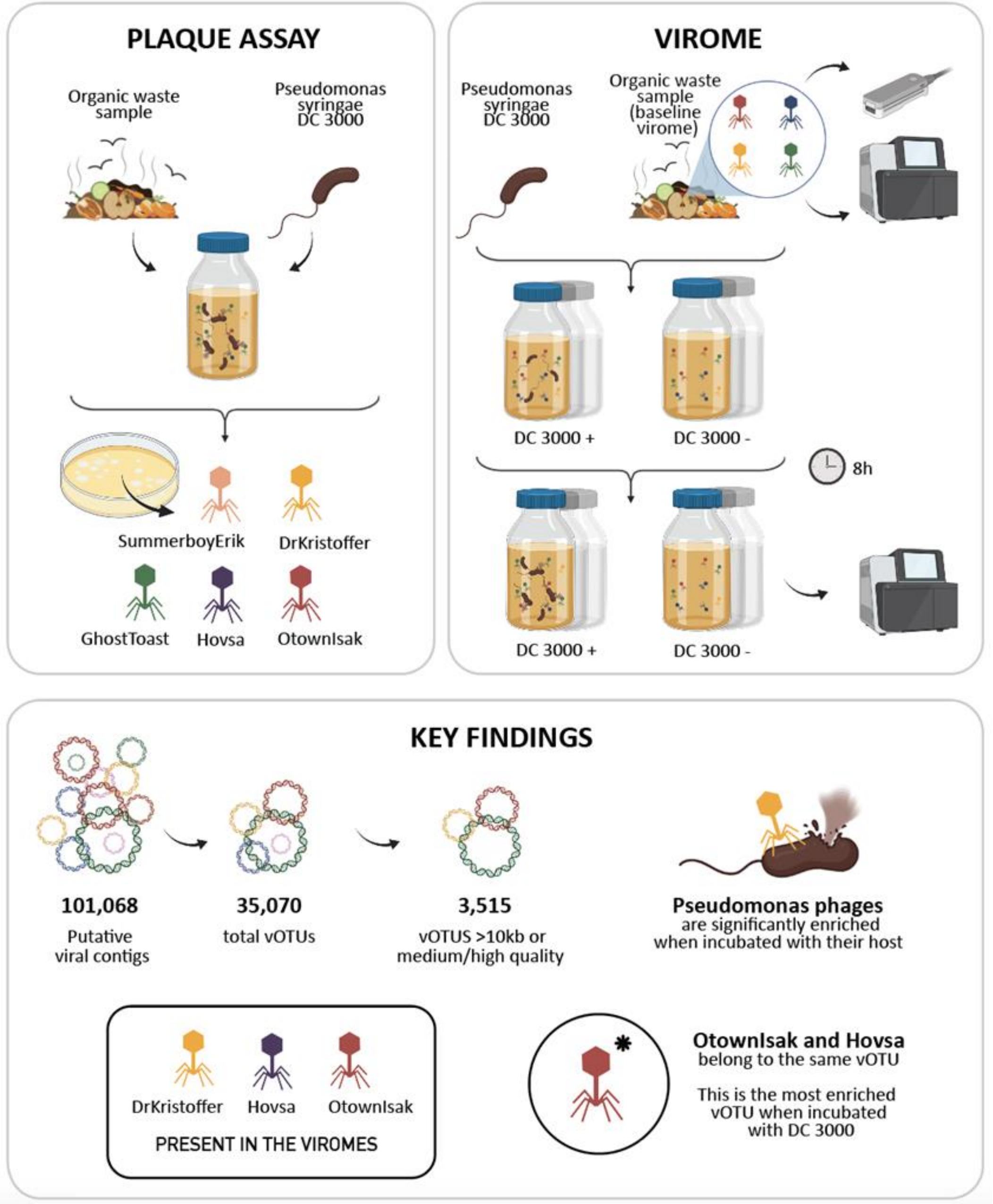
