## Supplementary material for "Metaviromes reveal the dynamics of Pseudomonas host-specific phages cultured and uncultured by plaque assay": Figure S6

### Table S6

**Table S6:** Viral OTUs composition of the population present in the three replicates of virome DC3000 + and DC3000 and the baseline virome. **A** Hierarchical clustering of vOTUs based on Jaccard distances. **B** Pairwise Bray-Curtis distances. **C** PCoA plot. **D** Comparison of treatment – incubation of OW in LB media at room temperature for 8 hours with or without an overnight culture of DC3000. ANOSIM test statistic = 0.42 (p-value = 0.106).

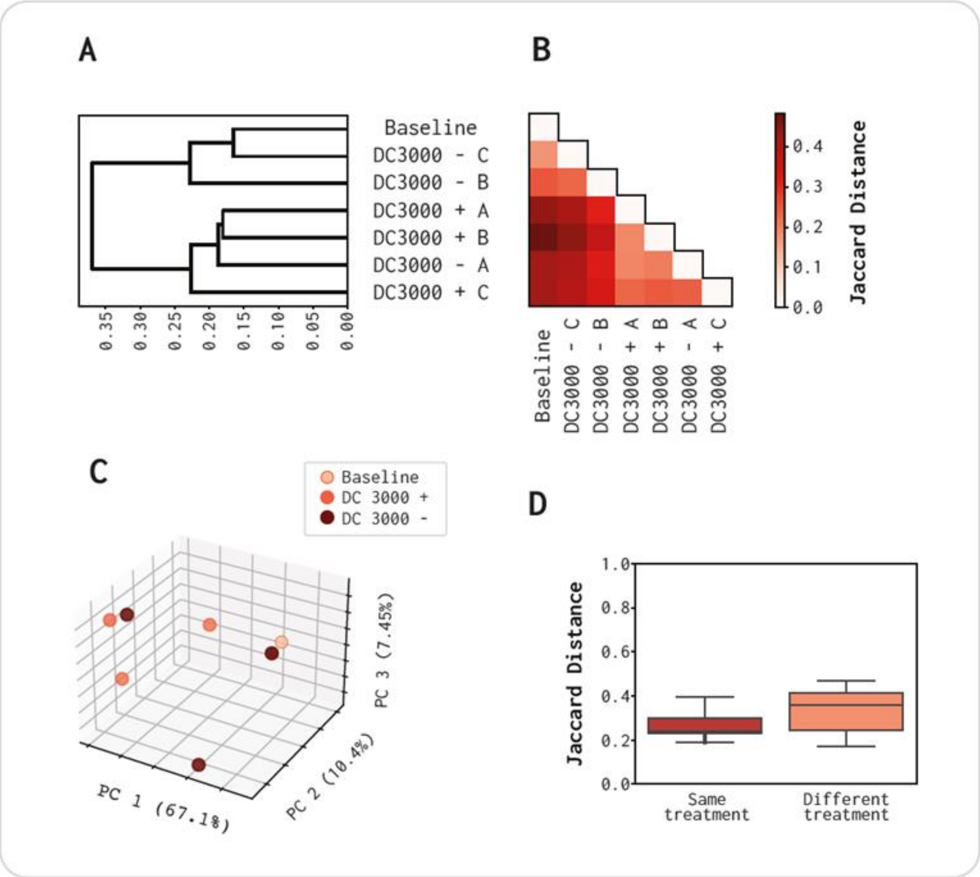
