## Supplementary figures and images for "Metaviromes reveal the dynamics of Pseudomonas host-specific phages cultured and uncultured by plaque assay"

### Figure S7

**Figure S7**

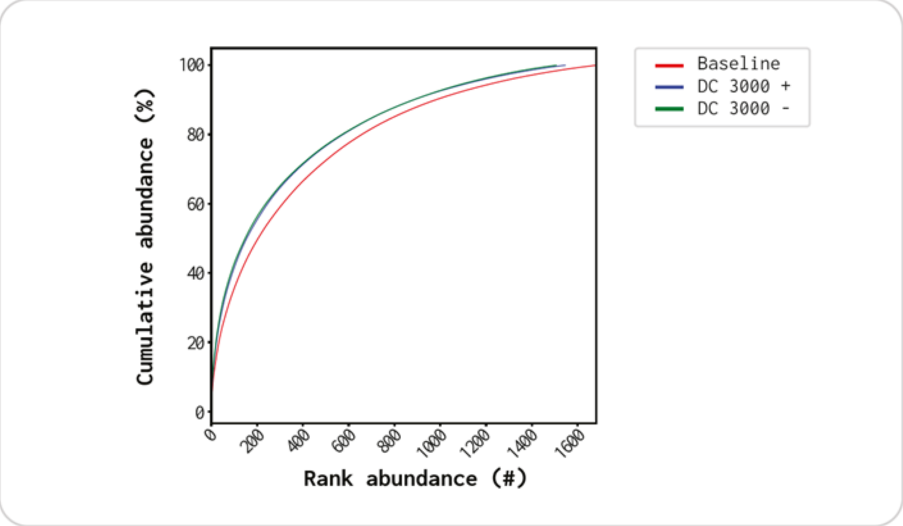


**Figure S7:** Accumulative abundance of all contigs in the three virome types.
