## Supplementary material for "Metaviromes reveal the dynamics of Pseudomonas host-specific phages cultured and uncultured by plaque assay": Table S2

**Table S2**: Overview of DNA amount, quality, and reads after quality control (QC) for each of the seven viromes used for the abundance analysis. N = Nanopore reads, P = paired Illumina reads, and UP = unpaired Illumina reads.

| **Virome** | **Total DNA (ng)** | **Nanodrop (260/280)/(260/230)** | **Post QC reads** |
|---|---|---|---|
| Baseline | 2,520 | (1.2) / (0.7) | N: 1251909, P: 9500926, UP: 453784 |
| DC3000+ A | 543 | (1.9) / (2.1) | P: 3641173, UP: 275037 |
| DC3000+ B | 474 | (1.8) / (2.1) | P: 4328342, UP: 350347 |
| DC3000+ C | 597 | (1.9) / (2.0) | P: 4558370, UP: 1861554 |
| DC3000- A | 4.76 | (0.3) / (0.2) | P: 5227108, UP: 333565 |
| DC3000- B | 25.6 | (0.4) / (0.3) | P: 4970642, UP: 438710 |
| DC3000- C | 24.4 | (0.4) / (0.3) | P: 5596835, UP: 408592 |
