## Supplementary material for "Metaviromes reveal the dynamics of Pseudomonas host-specific phages cultured and uncultured by plaque assay": Table S3

**Table S3:** Virsorter2 results on contigs. Data of total contigs, as well as how much was identified putative viral and putative non-viral by VirSorter2.

| Assembly | No. of putative viral contigs | No. of putative non-viral contigs | Total no. of contigs | % putative viral contigs | % putative non-viral contigs |
|---|---|---|---|---|---|
| Baseline metaSPAdes (Assembly 1) | 11,867 | 3,915 | 15,782 | 75.2 | 24.8 |
| DC3000 + metaSPAdes (Assembly 2) | 9,481 | 2,690 | 12,171 | 77.9 | 22.1 |
| DC3000 + A metaSPAdes (Assembly 3) | 3,738 | 883 | 4,621 | 80.9 | 19.1 |
| DC3000 + B metaSPAdes (Assembly 4) | 4,155 | 1,091 | 5,246 | 79,2 | 20.8 |
| DC3000 + C metaSPAdes (Assembly 5) | 2,818 | 686 | 3,504 | 80.4 | 19.6 |
| DC3000 – metaSPAdes (Assembly 6) | 16,925 | 5,332 | 22,257 | 76.0 | 24.0 |
| DC3000 – A metaSPAdes (Assembly 7) | 6,796 | 1,904 | 8,700 | 78.1 | 21.9 |
| DC3000 – B metaSPAdes (Assembly 8) | 6,389 | 1,860 | 8,249 | 77.5 | 22.5 |
| DC3000 – C metaSPAdes (Assembly 9) | 6,647 | 1,865 | 8,512 | 78.1 | 21.9 |
| All samples Hybrid metaSPAdes (Assembly 10) | 30,068 | 9,666 | 39,734 | 75.7 | 24.3 |
| Baseline Canu (Assembly 11) | 2,184 | 113 | 2,297 | 95.1 | 4.9 |
| Baseline Canu (Assembly 12) | 2,184 | 113 | 2,297 | 95.1 | 4.9 |
