## Supplementary material for "Metaviromes reveal the dynamics of Pseudomonas host-specific phages cultured and uncultured by plaque assay": Table S4

**Table S4:** Virsorter2 results on bp. Data of total bp, as well as how much was identified putative viral and putative non-viral by VirSorter2

| Assembly | No. of putative viral bp | No. of putative non-viral bp | Total no. of bp | % putative Viral bp | % putative non-viral bp |
|---|---|---|---|---|---|
| Baseline metaSPAdes (Assembly 1) | 29,835,217 | 5,643,379 | 35,478,596 | 84.1 | 15.9 |
| DC3000 + metaSPAdes (Assembly 2) | 21,504,440 | 3,661,585 | 25,166,025 | 85.5 | 14.5 |
| DC3000 + A metaSPAdes (Assembly 3) | 9,004,304 | 1,196,940 | 10,201,244 | 88.3 | 11.7 |
| DC3000 + B metaSPAdes (Assembly 4) | 9,500,589 | 1,466,266 | 10,966,855 | 86.6 | 13.4 |
| DC3000 + C metaSPAdes (Assembly 5) | 6,364,929 | 926,243 | 7,291,172 | 87.3 | 12.7 |
| DC3000 – metaSPAdes (Assembly 6) | 39,008,458 | 7,481,007 | 46,489,465 | 83.9 | 16.1 |
| DC3000 – A metaSPAdes (Assembly 7) | 15,563,712 | 2,596,593 | 18,160,305 | 85.7 | 14.3 |
| DC3000 – B metaSPAdes (Assembly 8) | 14,712,604 | 2,586,311 | 17,298,915 | 85.0 | 15.0 |
| DC3000 – C metaSPAdes (Assembly 9) | 15,557,533 | 2,593,590 | 18,151,123 | 85.7 | 14.3 |
| All samples hybrid metaSPAdes (Assembly 10) | 86,640,254 | 14,128,443 | 100,768,697 | 86.0 | 14.0 |
| Baseline Canu (Assembly 11) | 27,445,358 | 314,256 | 27,759,614 | 98.9 | 1.1 |
| Baseline Canu (Assembly 12) | 27,436,635 | 313,952 | 27,750,587 | 98.9 | 1.1 |
