## Supplementary material for "Metaviromes reveal the dynamics of Pseudomonas host-specific phages cultured and uncultured by plaque assay": Table S5

**Table S5:** CheckV result on individual assemblies. Distribution of contig quality identification by CheckV in regards to phage genome completeness for each of the 11 assemblies

| Assembly | Complete/high | Medium | Low | Not determined | Total vOTUs |
|---|---|---|---|---|---|
| Baseline metaSPAdes (Assembly 1) | 29 | 51 | 12,274 | 2,434 | 14,788 |
| DC3000 + metaSPAdes (Assembly 2) | 17 | 31 | 9,865 | 1,532 | 11,445 |
| DC3000 + A metaSPAdes (Assembly 3) | 6 | 19 | 3,805 | 492 | 4,916 |
| DC3000 + B metaSPAdes (Assembly 4) | 7 | 20 | 4,346 | 582 | 4,955 |
| DC3000 + C metaSPAdes (Assembly 5) | 5 | 11 | 2,941 | 385 | 3,342 |
| DC3000 – metaSPAdes (Assembly 6) | 26 | 52 | 17,504 | 3,204 | 20,786 |
| DC3000 – A metaSPAdes (Assembly 7) | 15 | 19 | 7,060 | 1,048 | 8,142 |
| DC3000 – B metaSPAdes (Assembly 8) | 9 | 28 | 6,808 | 899 | 7,744 |
| DC3000 – C metaSPAdes (Assembly 9) | 12 | 23 | 6,912 | 1,044 | 7,991 |
| All samples Hybrid metaSPAdes (Assembly 10) | 132 | 196 | 29,751 | 6,401 | 36,480 |
| Baseline Canu (Assembly 11) | 116 | 171 | 1,595 | 122 | 2,004 |
