## Supplementary material for "Metaviromes reveal the dynamics of Pseudomonas host-specific phages cultured and uncultured by plaque assay": Table S8

**Table S8:** Observed vOTUs in each of the virome samples used for Figure 4 in the main text.

| Sample | Observed vOTUs |
|---|---|
| Baseline | 1387 |
| DC3000 + A | 1547 |
| DC3000 + B | 1450 |
| DC3000 + C | 1316 |
| DC3000 – A | 1508 |
| DC3000 – B | 1500 |
| DC3000 – C | 1417 |
