## Supplementary material for "Metaviromes reveal the dynamics of Pseudomonas host-specific phages cultured and uncultured by plaque assay": Table S9

**Table S9:** Read mapping coverage of plaque-forming phages from organic waste sample. Total reads of each sample were mapped to the genomes. Depth coverage (black): trimmed mean pileup coverage. Breadth coverage (red) Percentage of the contig positions covered by mapped reads. Gray: genomes with breadth coverage >=80%.

| Phage | Viral population | Baseline depth | Baseline breadth | All depth | All breadth |
|---|---|---|---|---|---|
| DrKristoffer | NODE_5 | 0.103 | 51.28% | 3.929 | 98.86% |
| GhoastToast | GhoastToast | 0.023 | 25.49% | 0.018 | 34.55% |
| Hovsa | NODE_16 | 0.015 | 21.77% | 0.295 | 60.04% |
| OtownIsak | NODE_16 | 0.065 | 50.70% | 4.393 | 99.99% |
| SummerBoyErik | SummerBoyErik | 0.037 | 32.17% | 0.024 | 42.61% |

| Phage | Viral population | DC3000 + A depth | DC3000 + A breadth | DC3000 + B depth | DC3000 + B breadth | DC3000 + C depth | DC3000 + C breadth | DC3000 + depth | DC3000 + breadth |
|---|---|---|---|---|---|---|---|---|---|
| DrKristoffer | NODE_5 | 52.287 | 94.52% | 0.310 | 55.23% | 0.538 | 59.72% | 16.562 | 98.25% |
| GhoastToast | GhoastToast | 0.000 | 4.77% | 0.000 | 0.84% | 0.000 | 1.89% | 0.000 | 6.67% |
| Hovsa | NODE_16 | 0.529 | 37.02% | 1.141 | 43.88% | 0.972 | 42.21% | 0.937 | 49.57% |
| OtownIsak | NODE_16 | 14.643 | 99.14% | 18.284 | 99.12% | 11.018 | 99.06% | 15.151 | 99.99% |
| SummerBoyErik | SummerBoyErik | 0.000 | 2.21% | 0.000 | 2.80% | 0.000 | 1.24% | 0.000 | 5.30% |

| Phage | Viral population | DC3000 - A depth | DC3000 - A breadth | DC3000 - B depth | DC3000 - B breadth | DC3000 - C depth | DC3000 - C breadth | DC3000 - depth | DC3000 - breadth |
|---|---|---|---|---|---|---|---|---|---|
| DrKristoffer | NODE_5 | 0.083 | 35.36% | 0.053 | 26.80% | 0.062 | 31.47% | 0.081 | 54.39% |
| GhoastToast | GhoastToast | 0.000 | 6.69% | 0.013 | 15.06% | 0.010 | 14.36% | 0.014 | 23.90% |
| Hovsa | NODE_16 | 0.157 | 36.16% | 0.154 | 38.32% | 0.029 | 20.41% | 0.126 | 47.26% |
| OtownIsak | NODE_16 | 3.014 | 98.78% | 1.837 | 97.30% | 0.494 | 86.67% | 1.793 | 99.04% |
| SummerBoyErik | SummerBoyErik | 0.000 | 8.82% | 0.014 | 15.46% | 0.012 | 15.62% | 0.018 | 28.42% |
