## Supplementary material for "Metaviromes reveal the dynamics of Pseudomonas host-specific phages cultured and uncultured by plaque assay": Table S10

**Table S10:** CRISPR hits on host taxonomy at species level. Out of the 3,515 viral populations, 1164 had hits to a bacterial CRISPR spacer from the Shmakov database. A total of 806 of those hits could be assigned to a host at the species level. Below are the genera with ≥ 5 assigned viral populations.

| Genus | Count | Genus | Count |
|---|---|---|---|
| *Lactobacillus* | 209 | *Bifidobacterium* | 10 |
| *Pseudomonas* | 47 | *Bacillus* | 9 |
| *Xanthomonas* | 18 | *Salmonella* | 7 |
| *Dickeya* | 18 | *Aeromonas* | 7 |
| *Acinetobacter* | 17 | *Klebsiella* | 7 |
| *Pediococcus* | 14 | *Delftia* | 6 |
| *Streptococcus* | 13 | *Yersinia* | 6 |
| *Escherichia* | 12 | *Pluralibacter* | 5 |
| *Serratia* | 11 | *Porphyromonas* | 5 |
| *Acetobacter* | 11 | *Bacteroides* | 5 |
