## Supplementary material for "Metaviromes reveal the dynamics of Pseudomonas host-specific phages cultured and uncultured by plaque assay": Table S11

**Table S11:** Putative phage contigs that has a Log2FC above 1 (Green dots) or below – 1 (Red dots) (Figure 5A). Contigs were blasted using megablast. Denoted below are the top viral hit with lowest E-value and highest query coverage.

| Contig | Size (Bp) | > 1: Log2FC > 1<br>< -1: Log2FC < -1 | Blast hit (Accession no.) | Query Coverage | Per. Identity | E-Value |
|---|---|---|---|---|---|---|
| tig00010143 | 20829 | > 1 | Lactobacillus phage iLp1308 (KR905070.1) | 7% | 77.16% | 0.0 |
| NODE_45 | 50899 | > 1 | Siphoviridae NHS-Seq1 (MH029512.1) | 32% | 80% | 0.0 |
| tig00000588 | 18290 | > 1 | Lactobacillus phage 3-SAC12 (MK504442.1) | 3% | 91.75% | 2e-161 |
| NODE_3 | 59207 | > 1 | Enterobacteria phage CAjan (KP064094.1) | 100% | 100% | 0.0 |
| NODE_45 | 10380 | > 1 | Lactococcus phage 56301 (NC_049405.1) | 45% | 85.77% | 0.0 |
| NODE_220 | 5166 | > 1 | Lactobacillus phage AQ113 (HE956704.1) | 8% | 84.40% | 3e-119 |
| NODE_2743 | 5420 | > 1 | Leuconostoc phage CHB (KX578043.1) | 8% | 67.40% | 3e-29 |
| tig00001464 | 9726 | < -1 | Lactobacillus phage Lenus (NC_047897.1) | 89% | 97.35% | 0.0 |
| NODE_830 | 5136 | < -1 | Paramecium bursaria Chlerella virus CVM-1 (JX997163.1) | 1% | 75.29% | 0.003 |
| NODE_690 | 6178 | < -1 | Lactobacillus phage 3-521 (NC_048753.1) | 100% | 97.28% | 0.0 |
| NODE_2693 | 5480 | < -1 | Lactobacillus phage 3-521 (NC_048753.1) | 93% | 97.70% | 0.0 |
| NODE_132 | 40054 | < -1 | Uncultured Caudivirales phage (LR796317.1) | 1% | 68.32% | 1e-37 |
| tig00000648 | 32820 | < -1 | Lactobacillus phage LR2 (MH837543.1) | 5% | 67.97% | 3e-64 |
| tig00000040 | 44196 | < -1 | Escherichia phage C130_2 (NC_048067.1) | 2% | 76.66% | 4e-127 |
| NODE_852 | 11518 | < -1 | Phage apr34_1789 (MK415401.1) | 0% | 80.21% | 2e-11 |
| tig00000828 | 11199 | < -1 | Proteus phage vB_PmiP_RS8pmA (MG575419.1) | 88% | 93.04% | 0.0 |
| NODE_830 | 5580 | < -1 | *Lactococcus lactis. Subs. Lactis strain G423* (CP024958.1) | 1% | 80% | 0.047 |
| NODE_214 | 32484 | < -1 | Proteus phage PM75 (NC_027363.1) | 61% | 92.01% | 0.0 |
