## Supplementary material for "Metaviromes reveal the dynamics of Pseudomonas host-specific phages cultured and uncultured by plaque assay": Table S12

**Table S12:** Putative phage contigs that has a Log2FC between -1 and 1 without a significant abundance (Figure 5A). Contigs were blasted using megablast. Denoted below are the top viral hit with lowest E-value and highest query coverage.

| Contig | Size (Bp) | 0-1: Log2FC 0-1<br>-1-0: Log2FC -1-0 | Blast hit (accession no.) | Query coverage | Per. identity | E-value |
|---|---|---|---|---|---|---|
| NODE_364 | 6226 | 0-1 | Proteus phage vB_PmiP_RS51pmB (MG575421.1) | 6% | 79.59% | 5e-44 |
| NODE_209 | 8610 | 0-1 | Bacteriophage sp. Isolate 179 (MN855830.1) | 54% | 84.73% | 0.0 |
| NODE_445 | 5408 | 0-1 | Lactobacillus phage ATCCB (MK504445.1) | 21% | 76.79% | 1e-174 |
| tig00001258 | 5764 | 0-1 | Lactobacillus phage AQ113 (HE956704.1) | 16% | 72.53% | 1e-154 |
| NODE_2231 | 6180 | 0-1 | Myoviridae sp. ctThM1 (MN582070.1) | 9% | 72.78% | 9e-50 |
| NODE_507 | 5015 | 0-1 | Bacteriophage sp. Isolate 103 (MN855801.1) | 3% | 73.96% | 1e-15 |
| NODE_165 | 6157 | 0-1 | Bacteriophage sp. Isolate 32 (MN855637.1) | 0% | 88.89% | 4e-05 |
| NODE_281 | 9947 | -1-0 | Xylella phage Usme (LR743523.1) | 1% | 75.94% | 1e-46 |
| NODE_244 | 10727 | -1-0 | Salmonella phage assan (MT074440.1) | 20% | 69.49% | 3e-97 |
| tig00000616 | 17334 | -1-0 | Lactobacillus phage CL2 (KR905067.1) | 5% | 83.54% | 0.0 |
| tig00010057 | 16105 | -1-0 | Lactobacillus phage LBR48 (GU967410.1) | 8% | 92.67% | 0.0 |
| NODE_966 | 5065 | -1-0 | Bacteriophage sp. isolate 230 (MN855846.1) | 41% | 83.45% | 0.0 |
| NODE_1644 | 7563 | -1-0 | Lactobacillus phage ATCC 8014-B2 (NC_047739.1) | 22% | 80.31% | 0.0 |
| NODE_2505 | 5716 | -1-0 | Lactobacillus phage ATCC 8014-B2 (NC_047739.1) | 84% | 85.48% | 0.0 |
